# Opposing mechanisms of competition determine species invasions and functional diversity

**DOI:** 10.1101/2021.01.25.428088

**Authors:** Mark K. L. Wong, Roger H. Lee, Chi-Man Leong, Owen T. Lewis, Benoit Guénard

**Author notes:** Corresponding author: Mark K. L. Wong.

## Abstract

Understanding how species’ phenotypic differences affect competition is key to explaining community assembly and predicting biodiversity responses. Many studies overlook the variable effects that species’ trait differences can have on the direction of competitive exclusion, which reverses depending on the specific mechanism at play. We performed a comprehensive trait-based study of an ant invasion integrating morphological, dietary, physiological and behavioral analyses. We found that trait differences between invasive and resident species were not only associated with niche differences which promoted the coexistence of dissimilar species, but also competitive ability differences which acted in the opposite fashion. Furthermore, competition along separate trait axes led to complex and contrasting patterns in the invaded assemblages, where species were at once similar (clustered) in some traits but also dissimilar (overdispersed) in others. Our results reveal that different aspects of phenotype may distinctly modulate the effect of competition in structuring ecological communities and functional diversity.

## Introduction

Interspecific competition is a fundamental force structuring and maintaining biodiversity (1–3) and can also modulate the effects of climate change, habitat alteration and biological invasions (4–6). An essential goal in ecology is therefore to predict the outcome of interspecific competition on the basis of general and measurable properties of species, such as their functional traits (7). However, this is seldom achieved empirically because species’ trait differences can affect competitive outcomes in complex and contrasting ways.

Ecological theory suggests that trait differences can drive two contrasting effects on competitive outcomes between species (8–10). On the one hand, differences in species’ trait values may facilitate the use of different ecological niches, such as the exploitation of different resources, or activity at different times of the day. These ‘niche effects’ of trait differences tend to promote coexistence of species with dissimilar trait values and drive competitive exclusion among species with similar trait values [the concept of ‘limiting similarity’ (11)]. On the other hand, differences in trait values can also distinguish species in terms of their competitive abilities in a competition hierarchy, such as in the exploitation of a shared resource or activity period. In contrast to niche effects, these ‘competitive effects’ of trait differences promote coexistence among species with comparably strong competitive abilities and similar trait values, while driving the exclusion of other species with weaker competitive abilities and dissimilar trait values (8,12). Importantly, niche and competitive effects of trait differences generate contrasting patterns in the trait structures or ‘functional diversity’ of species assemblages: species tend to be overdispersed in assemblage trait space when niche effects are at play but clustered when competitive effects dominate (10,12).

Studies of plant assemblages have revealed the importance of both trait-based niche and competitive effects in determining coexistence outcomes and shaping community assembly. While high plant diversity has been attributed to interspecific niche differentiation across multiple trait axes (13), species’ differences in specific traits such as wood density, specific leaf area and maximum height have also been linked to competitive advantages that are consistent across multiple biomes globally (3). Notably, results from competition experiments suggest that both niche and competitive effects may operate simultaneously and associate variably with individual traits or across multidimensional trait space (14), and differently for morphological versus physiological traits (15). In most empirical studies of animals, however, limiting similarity is often assumed to be the sole basis for competitive exclusion. Interspecific trait variation is thus taken to reflect niche differentiation; and attempts to detect the effects of competition in community assembly typically involve tests for trait overdispersion only (16). The potential for trait differences to confer competitive advantages as opposed to niche differences may therefore be underestimated.

Invasions by non-native species provide an excellent opportunity to test mechanisms of community assembly (17). Trait differences of invasive and native plant species, such as leaf-area allocation and growth rate, have been shown to correspond to competitive advantages rather than niche differences (18), but the extent to which trait differences between invasive and native animal species promote niche and competitive differences, and the consequences for the functional structures of assemblages, are less well understood.

We investigated whether niche and competitive effects of trait differences in interspecific competition accounted for the outcomes of an invasion by the Red Imported Fire Ant, *Solenopsis invicta*, and their impacts on the functional structure of ground-foraging ant assemblages comprising 16 species in a tropical grassland. Using a novel comparative approach, we investigated how the presence and abundance of the invader *S. invicta* across 61 plots was explained by its trait differences with the ‘observed assemblage’ of ant species at each plot, and also with the ‘missing assemblage’ of ant species that were absent from each plot but present in the local species pool. Trait differences were measured in two ways: as absolute dissimilarities to approximate niche differences, and as hierarchical differences to approximate competitive differences [following (3,14)]. We investigated how the abundance of *S. invicta* related to its trait differences from missing species to understand competitive mechanisms potentially excluding ant species following invasion. Similarly, we scrutinized trait differences between *S. invicta* and observed assemblages to understand mechanisms promoting its abundance and potential coexistence with other ant species. Finally, we examined how interspecific competition with the invader *S. invicta* structures ant assemblages by comparing uninvaded and invaded assemblages in terms of the degree of overdispersion or clustering along individual trait axes as well as in multidimensional trait space.

Besides being among the first to assess both niche and competitive effects of trait differences in shaping an animal assemblage, our work goes further than previous studies by incorporating a diverse suite of traits spanning species’ morphology (seven traits), diet (Trophic Position), physiology (Critical Thermal Maximum), and behavior (Interference Ability) – all measured as continuous variables directly from multiple individuals of each species captured in the field. This work advances our understanding of how interspecific differences across the phenotypic spectrum drive competitive outcomes among species, and more broadly shows that trait differences can determine the responses of biodiversity to biotic disturbances.

## Results

The abundance of the invader *S. invicta* was well explained by its average trait differences with ant species in the missing assemblage at each plot (Fig. 1a-c) and those in the observed assemblage at each plot (Fig. 1d-f). Both niche and competitive effects were detected and associated with separate traits (Table 1; Fig. 1).

**Figure 1.**
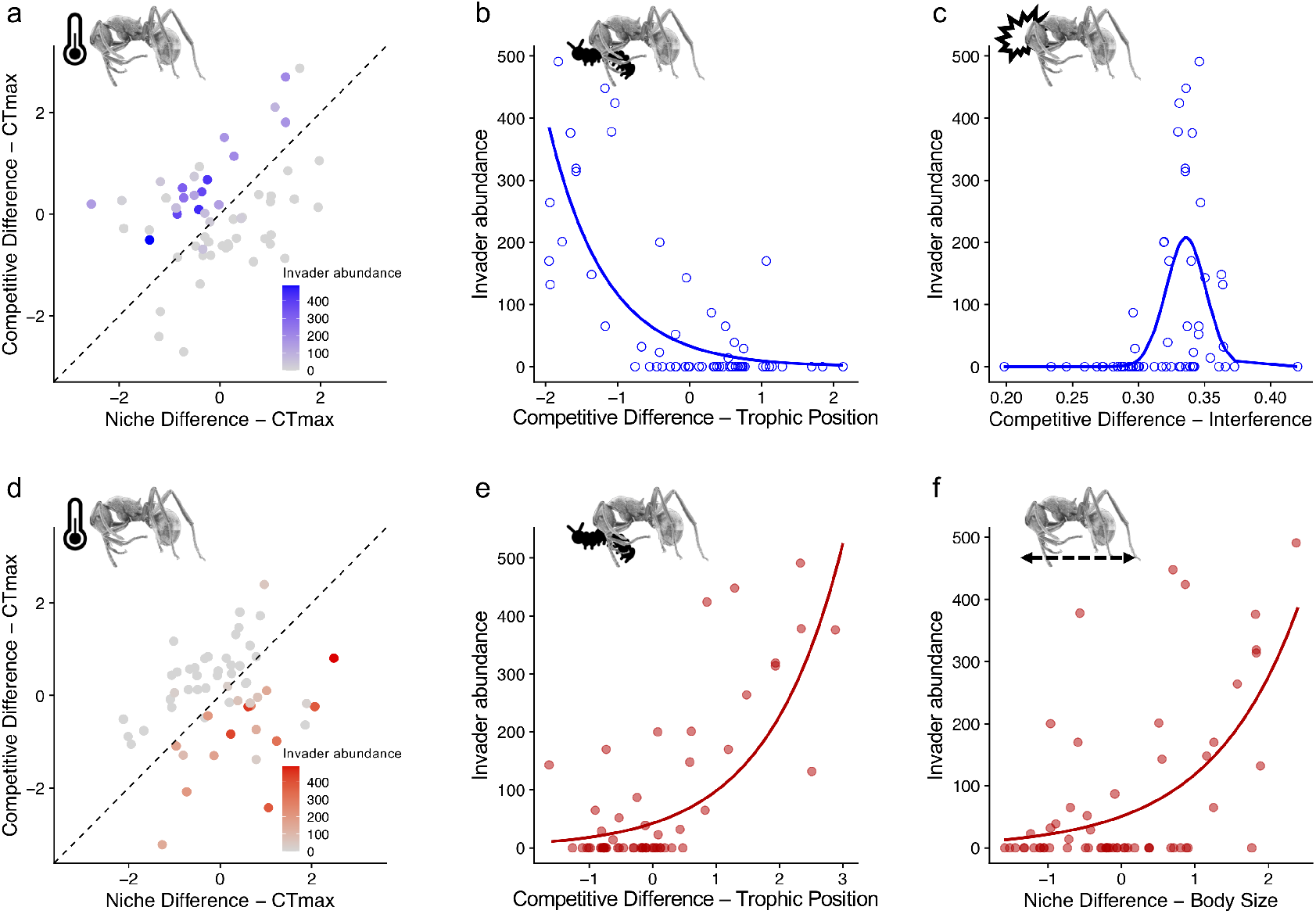
Abundance of the invasive species *S. invicta* as a function of its average trait differences with the ‘missing assemblage’ of ant species at each of the 61 plots (**a-c**), and with the ‘observed assemblage’ of ant species at each of the 61 plots (**d-f**). Against the missing assemblages, the model for Critical Thermal Maximum (CT_max_) detected significant effects of both Niche Difference and Competitive Difference, and predicted higher densities of *S. invicta* when Competitive Difference exceeded Niche Difference (**a**). The abundance of *S. invicta* was also significantly higher across plots where missing species tended to be more carnivorous than *S. invicta* (**b**), and where missing species had far poorer Interference Ability than *S. invicta* (**c**). Against the observed assemblages, the model for CT_max_ detected significant effects of both Niche Difference and Competitive Difference, and predicted higher abundances of *S. invicta* when Niche Difference exceeded Competitive Difference (**d**). The abundance of *S. invicta* was also significantly higher across plots where the species present were less carnivorous than *S. invicta* (**e**), and where the species present tended to be more dissimilar in Body Size (i.e., smaller or larger) from *S. invicta* (**f**).

**Table 1.** Results of trait and environment models for the abundance of the invasive ant *S. invicta* across 61 plots, with standardized coefficients. Trait models explain *S. invicta* abundance as a function of its average trait differences with either the ant species observed at each plot, or the ant species missing from each plot (but occurring in the species pool). Two types of trait differences between *S. invicta* and ant species were measured: Niche Difference and Competitive Difference. Environment models were built for the percentage ground cover (Ground Cover) and mean annual temperature (Temperature) at each plot. These variables were also included as covariates in trait models if they improved model performance.

| Model Type | Model Name | AICc | Terms | $\beta$ | P |
| --- | --- | --- | --- | --- | --- |
| *Trait Difference: Invader vs. Observed* |  |  |  |  |  |
|  | Head Width | 401.6 | Competitive Difference | 4.5 | <0.001*** |
|  |  |  | Ground Cover | 1.77 | <0.05* |
|  | CT<sub>max</sub> | 398.1 | Competitive Difference | -3.32 | <0.001*** |
|  |  |  | Niche Difference | 3.02 | <0.001*** |
|  | Trophic Position | 409.6 | Competitive Difference | 0.84 | <0.05* |
|  | Body Size | 411.7 | Niche Difference | 0.85 | <0.05* |
| *Trait Difference: Invader vs. Missing* |  |  |  |  |  |
|  | Head Width | 409.1 | Competitive Difference | -5.01 | <0.001*** |
|  |  |  | Ground Cover | 2.09 | <0.05* |
|  | CT<sub>max</sub> | 401 | Competitive Difference | 4.18 | <0.001*** |
|  |  |  | Niche Difference | -3.44 | <0.001*** |
|  | Interference | 396.9 | Competitive Difference | 3.1 | <0.001*** |
|  |  |  | Competitive Difference<sup>2</sup> | -3.17 | <0.001*** |
|  | Body Size | 424.2 | Niche Difference | -2.46 | <0.05* |
|  | Trophic Position | 405.9 | Competitive Difference | -1.29 | <0.001*** |
|  |  |  | Temperature | 0.62 | 0.11 |
| *Environment* |  |  |  |  |  |
|  | Ground Cover | 428.5 | Ground Cover | 0.2 | 0.79 |
|  | Temperature | 428.2 | Temperature | 0.51 | 0.54 |

Only differences in Body Size were consistently associated with niche effects (Table 1). Resident species of similar Body Size to *S. invicta* were largely absent from plots where the invasive ant was abundant; these plots instead contained resident species with dissimilar Body Size (Fig. 1f). Differences in Interference Ability, Trophic Position and Head Width were associated with strong competitive effects. Plots with higher abundances of *S. invicta* contained species of comparable Interference Ability to *S. invicta* and which had less carnivorous diets and relatively narrower heads than *S. invicta* (Table 1; Fig. 1b,c). Species missing from these plots had much lower (weaker) Interference Ability, more carnivorous diets, and relatively wider heads than *S. invicta* (Table 1; Fig. 1e).

Interestingly, differences in Critical Thermal Maximum (CT_max_) were associated with both niche and competitive effects. The invader *S. invicta* was more abundant at plots where the missing ant species had relatively lower CT_max_ or otherwise an absolutely similar CT_max_ to *S. invicta*, with the former driving a stronger effect (Table 1; Fig. 1a). In turn, ant species observed to occur with *S. invicta* at these plots had relatively higher CT_max_ or otherwise an absolutely dissimilar CT_max_ from *S. invicta*, with the former driving a stronger effect (Table 1; Fig. 1d).

Compared to uninvaded ant assemblages, invaded assemblages had marginally lower species richness (M_Uninvaded_=6.84±1.76, M_Invaded_=5.67±2.01, P=0.02) and were significantly more clustered in multidimensional trait space (Fig. 2a). Importantly, niche and competitive effects also structured the invaded assemblages along individual trait axes, with contrasting results. Consistent with competitive effects, invaded assemblages were significantly clustered in Interference Ability and Head Width relative to uninvaded assemblages (Fig. 2b,d). However, consistent with niche effects, invaded assemblages were more overdispersed in CT_max_ than uninvaded assemblages (Fig. 2c).

**Figure 2.**
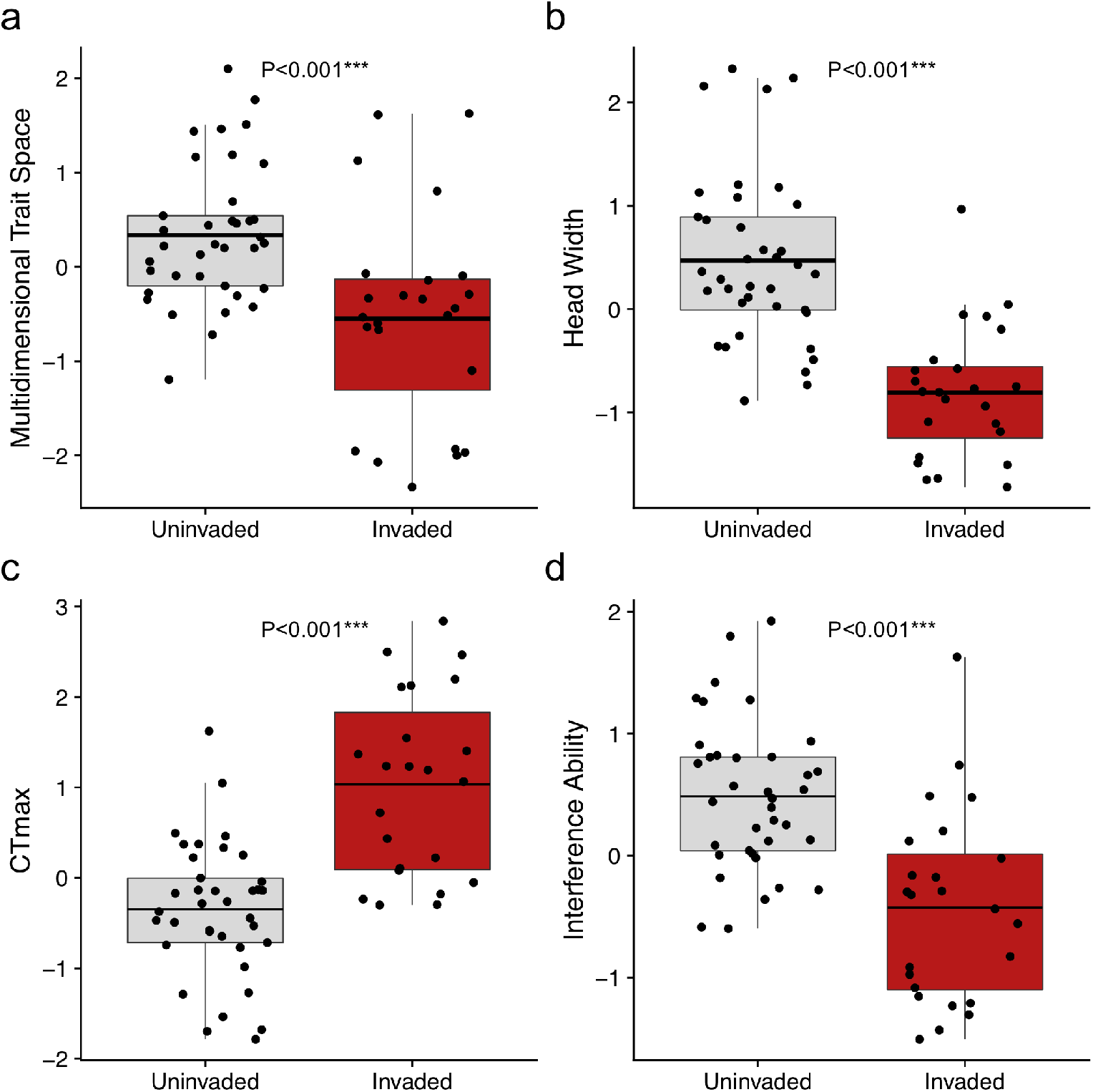
Standardized Effect Sizes (SES) for functional dispersion in Multidimensional Trait Space (**a**) and three separate traits – Head Width (**b**), Critical Thermal Maximum (CT_max_) (**c**), and Interference Ability (**d**) – across 37 uninvaded and 24 invaded ground-foraging ant assemblages. In comparison with the uninvaded assemblages, ants in the invaded assemblages were significantly more overdispersed in CT_max_ but significantly more clustered in Multidimensional Trait Space, Head Width and Interference Ability. P-values correspond to results from t-tests (**b**,**c**) and Wilcoxon Mann-Whitney tests (**a**,**d**). The invader *S. invicta* is excluded from the invaded assemblages in these analyses. Full details in Supporting information.

As expected from the study design, there was little evidence to suggest that environmental effects influenced invasion outcomes across the 61 plots (Table 1) as well as the species richness and functional structure of the ant assemblages (Supporting information).

## Discussion

A sound empirical understanding of how trait differences shape interspecific competition is fundamental to explaining community assembly, as well as for predicting biodiversity responses to environmental changes such as biological invasions. Here, we found that interspecific trait differences largely determined the outcomes of an invasion through opposing mechanisms of competitive exclusion (12). Specifically, differences between the invader *S. invicta* and resident ant species in separate morphological, dietary, physiological and behavioral traits facilitated two distinct mechanisms: limiting similarity driven by niche effects (11) and competitive hierarchies shaped by competitive effects (3,8,19). These mechanisms provide an explanation for the absence of certain species in invaded plots, and for the presence of others. Crucially, the two mechanisms acted distinctly on separate traits, and their opposing effects generated contrasting structures of overdispersion and clustering among those traits in the invaded assemblages. Our findings cast light on the thus far underestimated role of traits in conferring competitive advantages (in addition to niche opportunities) to species, underscoring their importance to interspecific competition in natural settings. The findings also illustrate how multiple, even opposing assembly mechanisms can simultaneously structure functional diversity.

A view of competition and coexistence, that extends beyond classical niche theory to incorporate stabilizing and equalizing mechanisms (8), is increasingly well-established in studies of plant assemblages (3,14). Our results suggest that such a perspective is equally relevant to understanding competition and coexistence in animals. Consistent with limiting similarity, niche differences led to the exclusion of species with similar body sizes to the invader *S. invicta*, while promoting coexistence between the invader and species that were sufficiently larger or smaller – a pattern observed in biological invasions of ants and a variety of taxonomic groups (20–22). However, we also found that niche differences alone did not determine competitive outcomes in the invasion; their stabilizing effects more likely counteracted the demographic effects of species’ differences in competitive abilities that would otherwise promote exclusion (8,9).

Contrary to the commonly held assumption that trait differences mainly reflect niche partitioning (12), we detected competitive effects associated with multiple traits of ant species. For instance, species with relatively wider heads and more carnivorous diets than *S. invicta* appeared to be excluded where *S. invicta* occurred, while species with relatively narrower heads and more carbohydrate-rich diets than *S. invicta* were observed in its presence. This may have occurred if *S. invicta* depleted arthropod prey populations, impacting more carnivorous ant species [as observed previously; (23)], driving a competition hierarchy in omnivory among native species. Another trait associated with strong competitive effects, as revealed from asymmetrical patterns of exclusion (Fig. 1c) and consequent clustering in assemblage trait structure (Fig. 2d), was interference ability. It has been proposed that interspecific trade-offs in stress tolerance and competitive dominance akin to those in plants (24) may likewise facilitate coexistence in ant assemblages (25). Given that *S. invicta* had very strong interference ability, and that other ant species in invaded assemblages had dissimilar or higher thermal tolerances (CT_max_) than *S. invicta* (Fig. 1d), such dominance-tolerance trade-offs may have been at play. Further supporting this hypothesis, ant species with higher thermal tolerances but lower interference ability than *S. invicta* were observed recruiting to baits abandoned by the invasive ant when those baits were heated by sun exposure following changes in cloud cover (Supporting information). Such trade-offs would represent equalizing mechanisms which, in addition to stabilizing mechanisms, promote coexistence between the invader and other ant species by mitigating interspecific differences in competitive abilities (8).

Our findings shed light on potential trait-based stabilizing and equalizing mechanisms that may determine the outcomes of competition among ant species. Rigorous tests for these coexistence mechanisms would require competition experiments measuring demographic parameters such as invasion growth rates (e.g. 14); such approaches are not readily transferable to animals (see 26). Nonetheless, we suggest that the potential for trait differences to reflect niche or competitive effects warrants explicit consideration of both in the structuring of empirical assemblages. We therefore agree with recent calls for community assembly studies to test for patterns of clustering in addition to overdispersion as signatures of competition (10,16). However, our findings also reveal important limitations in inferring community assembly processes exclusively from patterns in multidimensional trait space.

Of key relevance to observational studies, our results show that inferences about underlying ecological processes from patterns in the multidimensional trait structure of assemblages may be misguided if distinct processes with opposing effects act on different traits simultaneously. Here, potentially important effects of niche partitioning in body size (Fig. 1f) and thermal tolerances (Fig. 2c) in structuring the invaded assemblages were not well reflected in their strongly clustered multidimensional trait space (Fig. 2a). This was possibly due to the overwhelming, opposing effects of hierarchical competition along other trait axes such as head width (Fig. 2b) and interference ability (Fig. 2d). Such effects from multiple interactive and simultaneous assembly processes on functional structure are likely underestimated, especially where biotic interactions are concerned [but see (27)]. Yet these distinct competitive mechanisms have significant implications that reach beyond the fundamental understanding of community assembly. For instance, they can strongly determine biodiversity-ecosystem function relationships (28). While niche differences and resource partitioning drive complementarity effects, competitive hierarchies drive selection effects (28); each leads to a very different relationship between functional diversity and ecosystem functions (see 29).

Overall, our study uncovered new insights into the drivers and consequences of an ant invasion, which were consistent with the predictions of modern coexistence theory (8,12). Distinguishing between the contributions of traits to niche differences and competitive differences between species revealed alternative paths to competitive exclusion by invasion (30), and potential trait-based stabilizing and equalizing mechanisms structuring invaded assemblages. These specific mechanisms are useful for understanding invasion responses and assembly processes at fine spatial scales. The broader implication for empirical work is to embrace the potential for different traits across the phenotypic spectrum to reflect the varying role and effect of competition in structuring biodiversity.

## Materials and Methods

### Sampling ant assemblages and environmental variables

To minimize the effects of environmental and spatial processes on community assembly we focused on a simple grassland setting with low environmental heterogeneity and sampled at fine spatial scales well within ant species’ dispersal ranges. The study was conducted in two reserves of open grassland in Hong Kong which have been protected for >35 years, and which contain networks of exposed grass bunds (width ≤5 m) separating permanent ponds (31). Pilot surveys from 2015 to 2017 recorded colonies of *S. invicta* present at high densities at multiple locations (31). From April to September 2018, we systematically characterized the ground-foraging ant assemblages at 61 plots (each a 4 x 4 m quadrat; ≥25 m between adjacent plots) using pitfall traps followed by observations at baits. Sampling at this fine spatial scale allowed us to detect patterns driven by biotic interactions, since *S. invicta* and most ant species in the Oriental region forage within 5 m of their nests (32,33). For the same reasons, the minimum distance of 25 m between adjacent plots facilitated independent observations. The maximum distance between any two plots was 4 km; at this spatial scale, the effects of dispersal limitation were very likely minimized as all species disperse via flying alates which can reach far greater distances (34).

In each plot, six pitfall traps (diameter: 5.5 cm) were installed to sample the ants over a period of 48 hrs. Baits were then installed within 72 hrs from the retrieval of the pitfall traps. Between 1000-1500 hrs, five bait stations were installed in each plot, each comprising a slice of chicken sausage (diameter: 20 mm; height: 2 mm) on a white plastic disc (diameter: 5 cm) flushed with the ground. The sausage bait contained trophic resources required by most ants: proteins, lipids, carbohydrates and sodium (35). Each baiting session lasted 40 min, during which photographs were taken with a digital camera at 5 min intervals; these were subsequently used to determine the species recruiting, their abundances, and interspecific interactions (see ‘Behavioral trait: Interference Ability’ below). The mean ground surface temperature around each bait during the 40 min session was recorded with a thermogun. In pilot trials, the 40-min duration allowed for competitive interactions to reach unequivocal outcomes and for baits to be monopolized by single species. After each baiting session, live workers of all species encountered were collected and used for dietary and physiological trait measurements in the laboratory (below). Specimens collected from pitfall traps were used for morphological measurements, confirming species identities, and determining the occurrences of species in plots. While several species collected in pitfall traps were not recorded at baits (but not vice versa), many of these were hypogaeic. Given that our study aimed to investigate competition, we focused the analyses on the pool of species recorded at baits, as these best represented the ground-foraging ant assemblage limited by common resources.

At each plot, we also estimated the percentage of ground cover (Ground Cover) by applying color thresholding techniques in ImageJ (36) to digital photographs, and obtained high resolution (30 x 30 m) estimates of mean annual temperature (Temperature) from local climate models (37). As these environmental factors were shown to influence ant diversity in other invaded systems (e.g. 38) we used our data to test for environmental effects on invasion outcomes (the abundance of *S. invicta* across the plots), species richness and the trait structures of the assemblages.

### Morphological traits

We measured seven morphological traits (Body Size, Head Width, Eye Width, Mandible Length, Scape Length, Pronotum Width, Leg Length) on ≥10 individual workers of every species (N=319). These traits are linked to ant physiology and behavior and are hypothesized to impact performance and fitness (Supporting information).

### Dietary trait: Trophic Position

We measured the relative Trophic Position of each ant species using stable isotope ratios of Nitrogen (δ15N) (39). Live ants collected from the field were killed in a −20 °C freezer. We then rinsed the ants with distilled water, removed their abdomens to avoid contamination by undigested material in the gut (39), and dried the samples in an oven at 40 °C until a constant mass was reached. Dried samples with ≥5 workers were transferred into an aluminum capsule weighing 0.3–1 mg [workers of larger species were first ground and homogenized using a mortar and pestle following (40)]. We measured the δ15N values of each sample using a Nu Perspective Isotope Ratio Mass Spectrometer coupled to an Eurovector Elemental and reported in ‰ (41). Mineral soil collected from the field was used for baseline calibration. The δ15N value of every species was determined using 1–3 samples.

### Physiological trait: Critical Thermal Maximum

We measured the Critical Thermal Maximum (CT_max_) of individual ants following established protocols for CT_max_ assays (42). The ants were first acclimated at 25 °C for ≥2 hrs in the laboratory. Individual ants were then placed in 1.5 mL Eppendorf tubes. The entrance of each tube was plugged with dry cotton wool, ensuring that each ant was confined to an area of even temperature distribution. The tubes were then placed in a digital dry bath (BSH1004) connected to an additional thermometer (UEi Test Instruments DT302 Dual Input IP67) to ensure temperature accuracy. The assay began at a starting temperature of 36 °C, and the temperature was increased at a constant rate of 1 °C min^-1^ (42). Every 1 min, each tube was rotated and visually inspected to determine whether the ant had lost muscle coordination (42); the temperature at which this occurred was recorded as the individual’s CT_max_. We measured the CT_max_ of ≥10 individual workers of each species (N=193).

### Behavioral trait: Interference Ability

We assessed the Interference Ability of ant species from observations of interspecific interactions at baits after (43); therein termed ‘behavioral dominance’. Here, Interference Ability describes a species’ relative success in two types of antagonistic interactions: usurping a resource from heterospecifics (expulsion) and defending an occupied resource from usurping heterospecifics (retention) (43). We recorded the outcomes (win/loss) in expulsion and retention incidents between ant species at baits. Each species’ Interference Ability was then scored using the Colley rating method (43), which adjusted the value of each win and loss by the Interference Ability of the competitors that a species interacted with.

### Data analysis

#### Trait selection

For all morphological traits except Body Size, we corrected for the effects of body size by regressing each trait against Body Size and using the residuals as the new values for that trait. We used Principal Components Analysis (PCA) and correlation analysis to select a suite of traits that captured most interspecific variation in multidimensional trait space while minimizing redundancy from trait correlations (Supporting information). The first principal component, capturing 30% of the variation, was strongly positively associated with Head Width and Mandible Length and strongly negatively associated with Leg Length and Scape Length. The second (28% of the variation) was strongly positively and negatively associated with Interference Ability and Eye Width, respectively. The third (19% of the variation) was strongly positively and negatively associated with Body Size and Trophic Position, respectively. The fourth (13% of the variation) was strongly negatively associated with CT_max_. All subsequent principal components had eigenvalues lesser than unity. Among traits showing strong positive correlations (Supporting information), we selected those with stronger loadings on principal components. Our final set of traits comprised four morphological, one dietary, one physiological and one behavioral trait: Body Size, Head Width, Eye Width, Leg Length, Trophic Position, CT_max_, and Interference Ability.

#### Quantifying two measures of trait differences

For each trait, we quantified two measures of trait differences between the trait value of *S. invicta* (*T_S. invicta_*) and the trait value of every other ant species (*T_other_*) recorded at baits in the study. We calculated Niche Difference as |*T_S. invicta_ – T_Other_*|, a non-directional measure that can indicate the magnitude of niche differences between species. We calculated Competitive Difference as *T_S. invicta_ – T_other_*, a directional measure that detects differences in competitive abilities along a competition hierarchy (3,14).

#### Modelling invader abundance as a function of its trait differences with two assemblage types

The total number of *S. invicta* workers collected across the six pitfall traps at each of the 61 plots was used as the response term ‘Invader Abundance’. For each trait, we modelled Invader Abundance as a function of the average trait difference (including Niche Difference and Competitive Difference as separate terms) between *S. invicta* and the observed assemblage of ant species at each plot. We repeated this process using the average trait difference between *S. invicta* and the ‘missing assemblage’ at each plot, which comprised any species missing from the plot but occurring in the species pool. We modelled the relationships using Poisson models with observation-level random effects as well as negative binomial models (prior to selecting the best model), as both of these could address the overdispersion in Invader Abundance (44). For each trait, we built a full model which included Niche Difference, Competitive Difference, the quadratic form of Competitive Difference, and the environmental covariates Ground Cover and Temperature. We then selected the best model using a backward-stepwise variable selection procedure based on the Akaike information criterion corrected for small sample size (AICc).

#### Comparing patterns of trait dispersion between uninvaded and invaded assemblages

To determine whether invaded assemblages were more overdispersed or clustered relative to the uninvaded assemblages, we assessed the functional dispersion (FDisp) in each trait as well as in multidimensional trait space at the plot level (excluding *S. invicta*). We calculated FDisp using the ‘fdisp’ function of the FD package in R (45), and the ant species’ frequencies of occurrence across the six pitfall traps at each plot. To control for potential effects of species richness on FDisp, we compared standardized effect sizes (SES) instead of the observed values (46). We calculated SES by comparing the observed values to values generated from 999 constrained null models randomizing the matrix of species’ frequencies of occurrence using the ‘Independent Swap’ algorithm. The formula for calculating SES is:

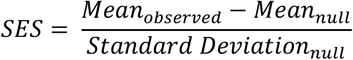

We then compared the SES values of FDisp in invaded plots to those in uninvaded plots using t-tests or Wilcoxon-Mann-Whitney tests (when sample variances were unequal).

## Acknowledgments

This study was supported by a National Geographic Grant (60–16) and a Clarendon Scholarship to MKLW, and an Environment and Conservation Fund Grant (32/2015) to BG. The authors are grateful to MTR and AEC for access to field locations, Sum Leung Kit for assistance with stable isotope measurements, Kaya Jumbe for assistance with baiting data, as well as Francois Brassard, Toby Tsang, Catherine Parr and Michelle Jackson for comments on previous versions of the manuscript.

